# Modelling the persistence of post-management disturbance in *Calluna vulgaris* communities

**DOI:** 10.64898/2026.05.12.724511

**Authors:** Jonathan Ritson, Benjamin Bell, Fred Worrall, Martin Evans, Richard Lindsay, Chris Evans

## Abstract

1. *Calluna vulgaris* is often managed in the UK by rotational burning, but this practice has recently been banned on peat with depth greater than 30-40 cm. It is unclear how then to manage the large areas of *Calluna* on blanket bogs used for sport shooting because without managed burning, fuel loads and wildfire risk will increase as the *Calluna* ages within the artificially narrow age distributions created by burn management.
2. We developed a model of *Calluna* mortality and management to understand duration and persistence of post-management effects. This allows us to assess how long it will take to reach a more natural age structure which would allow increased diversity if management ceases.
3. Our results show that management effects persist for around 50 years depending on site-specific mortality rates. Active management may therefore be needed either to mitigate the elevated risk of severe wildfire or to speed up this transition.
4. Some studies have employed, as unmanaged analogues, *Calluna* stands that were last managed <50 years ago, but such studies may have unintentionally biased their results by observing *Calluna* still in post-management recovery leading to an over-estimation of wildfire risk associated with more natural blanket bogs.
5. *Synthesis and applications*: with the banning of burning as a management tool for *Calluna* on deep peat, alternative management is now likely needed as our model shows it could take around 50 years for the *Calluna* to reach a more natural age distribution. Mowing can replicate some of the effects of managed burning but requires repeated intervention and may compress the peat surface from repeated machine tracking. Rewetting and *Sphagnum* reintroduction may offer a more sustainable management approach to lowering *Calluna* fuel loads and reducing severe wildfire risk by creating wetter sub-optimal conditions for *Calluna* growth and thereby altering the competitive balance between Sphagnum and *Calluna*. Further work is needed to assess the efficacy of rewetting in controlling fuel loads and how this varies with climate and local pressures. More broadly, this work highlights the need to quantify the persistence of past management regimes to understand ecological trajectories.

## Introduction

Peatland vegetation in the UK has been managed using fire in order to improve grazing conditions for centuries including to aid the removal of natural tree cover in the uplands (Davies et al., 2016). Use of managed fire increased with the process of enclosure, a legislative mechanism where many UK uplands were removed from a common ownership that came with self-negotiated rights and responsibilities amongst residents, to be replaced by private ownership and investment (Ritson and Lindsay, 2023). This change in management was often accompanied by increased liming and drainage to lower the water table and raise the pH of the peat, making the land more suitable for agriculture (Rennie et al., 1794). These processes, combined with the diffuse effects of pollution from the industrial revolution, led to widespread changes in vegetation across upland landscapes, particularly a general expansion of *Calluna vulgaris* (Holden et al., 2007) and the creation of the cultural landscapes seen on much of UK blanket bogs today. Whilst originally these management practices were undertaken to increase the capacity for sheep grazing, throughout the 19th century the rise in popularity of sport shooting of grouse (*Lagopus scotica*) led to a focus on increasing grouse numbers by promoting *Calluna* dominance as the significant driver of management in many UK upland environments (Moss 1904), though it should be noted that *Calluna* has dominated in many UK blanket bogs for hundreds of years at a time in the palaeoecological record (Chambers 1982, Chamber 1983, Chambers 2022, Webb et al. 2022, McCarroll et al. 2016)

The commonly used description *Calluna* growth strategy was given by Watt (1955), who described a 30-year life cycle divided into pioneer, building, mature and degenerate phases. In the pioneer phase, the young *Calluna* does not form a closed canopy and is interspersed with other plants. In the building phase the canopy closes, excluding nearly all light to the understory. In the mature phase, growth slows and gaps in the canopy form, before *Calluna* becomes ‘degenerate’, slowly collapsing and dying. The goal of management of *Calluna* has often been to truncate this lifecycle, preventing plants from reaching the degenerate stage and continuing *Calluna* dominance in perpetuity (Leslie, 1911).

Grouse management favours a patchwork of differing even-aged *Calluna* stands which provide both nesting cover and young shoots for food. This has often been achieved by burning patches of *Calluna* on a 15-to-25-year cycle. Early land managers favoured ‘back burning’, where the fire moves slowly and removes all vegetation (Leslie, 1911), whereas more recently ‘cool burns’ have been promoted as a means of burning only the *Calluna* canopy and leaving the bryophyte understory intact (Heather and Grass Burning Code (Defra, 2007)). Even though this modified form of burning management aims to preserve the understory vegetation, it has been deemed as potentially damaging to the peat (Noble et al. 2025). Consequently the UK Government has now banned burning on peat 30 cm deep or more in England, while the Scottish Government will, from autumn 2026, no longer permit burning on peat of 40+ cm depth for the purposes of sport - *i.e*. grouse shooting (Defra 2025; Wildlife Management and Muirburn (Scotland) Act 2024). This means that while managed burning can still be used on heaths with shallow organic soils, it will be banned on blanket bogs, the focus of our study, where it had previously been employed in the UK.

The potential cessation of burning as a management practice across much of the UK blanket bogs raises an important question about how land will be managed in the future if sheep grazing and grouse shooting continue to be the dominant economic drivers, particularly as ageing Calluna stands are associated with increased fuel loads and consequent wildfire risk (Allen et al. 2013). Options include the replacement of burning with mowing of *Calluna*, management away from *Calluna* dominance through rewetting and Sphagnum reintroduction, or the complete cessation of management in a ‘rewilding’ scenario.

If the land is to be kept in its current economic use (sheep grazing and grouse shooting) by land managers, each of these potential management options has advantages and disadvantages. However, given the extent and history of current *Calluna* management, it is difficult to assess implications of each, given the timescales required to understand an anthropogenic ecosystem currently managed on an approximately 15 to 25-year cycle. One approach currently employed is the use of chronosequences of *Calluna* of different ages at the same site as a proxy of what unmanaged *Calluna* might look like (Davies and Legg, 2008, Harris et al. 2011, Heinemeyer et al. 2023). A significant weakness with this approach is that it is unclear how long post-management disturbance of *Calluna* persists, which therefore raises the question of whether these sites can be used as true analogues of how the ecosystem would function in an unmanaged condition. Specifically, we do not know what the end point is for unmanaged *Calluna*, nor indeed how long it might take to reach a return to an unmanaged steady-state. Management by fire or cutting creates *Calluna* stands of a narrow age distribution. This means that canopy closure across a given stand happens uniformly, giving *Calluna* a strong competitive advantage and leading to lower vegetation diversity when the *Calluna* canopy closes (Gimingham, 1972; Watt, 1955). Conversely, a *Calluna* stand in a natural state should have plants in different age categories and different extents of canopy closure because young and old plants would be interspersed (Watt, 1955). It is important in assessing existing evidence, therefore, to establish whether previous studies have observed a truly unmanaged condition or simply observed a later stage in post-management disturbance.

An alternative approach to informing a better understanding of management implications for *Calluna* involves employing models of *Calluna* age structure. Previous attempts at modelling *Calluna* focused on fuel-load accumulation (i.e. above-ground woody biomass) and the interaction between management and wildfires (Allen et al. 2013). However, this model did not consider the effects on the population if *Calluna* were to be released from management. Furthermore, Allen et al., (2013) did not detail how they considered mortality in *Calluna* other than by fire, so they were unable to assess realistic age distributions in long-burn rotations. For example, in their 50-year return period, Allen et al.’s model shows ∼30% of plants aged 50 years despite this age not being observed in field observation studies (Forrest, 1971).

To explore *Calluna* age structure under different management scenarios on blanket bog and assess the validity of previous work using chronosequences, we therefore sought to develop a model which considers natural *Calluna* mortality as well as management. This allows us to generate more realistic age distributions within managed and unmanaged populations and thereby estimate how long post-management disturbance persists and what age distributions might look like under different management options. Such a model can shed a valuable light on previous studies which have used *Calluna* stands at set times post-management as a proxy for an unmanaged condition. Furthermore, using this model we can also assess how long *Calluna* would take to reach a stable state in a rewilding scenario and therefore estimate over what timescale both fuel-load and severe wildfire risk would likely be increasing in the absence of other interventions.

Our aim was therefore to a) to estimate the time taken to reach an unmanaged state after both catastrophic wildfire and the cessation of management, and b) assess if previous work using chronosequences of *Calluna* post-management were valid representations of an unmanaged state.

## Materials and Methods

Here we present a model of *Calluna* population ageing that offers a means of assessing the timescales required to move from a post-burn narrow distribution of age stand to a mixed-age stand that could be considered to be ‘unmanaged’

Our model assumes a) that post-disturbance, such as fire or cutting on an area that was previously dominated by *Calluna*, recovers rapidly through vegetative regrowth from stems and/or seedlings; b) *Calluna* plants age and die based on the age-dependent mortality rates given by Forrest (1971); c) that each dead *Calluna* plant is soon replaced by another living *Calluna* plant; and d) this cycle repeats.

Assumption (a) is widely accepted, as this is the stated aim of *Calluna* management by fire and cutting on grouse estates where cutting is used to mimic the effects of burning rather than for conservation. Assumption (b), we believe, gives a more accurate representation of reality than previous models of *Calluna* age-distribution (Allen et al., 2013) that used limited mortality effects and therefore had unrealistic age distributions for long burn rotations. However, our mortality rate is dependent on age only, and does not account for environmental or climatic effects, which could further influence mortality rates (Armstrong et al., 1997).

Whilst assumption (c) is possible (see Barclay-Estrup and Gimingham, 1969; Gimingham, 1972 for discussion and photographs of pioneer *Calluna* emerging from degenerate *Calluna*), there can often be potentially long periods (Gimingham, 1972) of either bare peat or periods where other vascular plants encroach before *Calluna* can re-establish itself. Our model can replicate this effect by including a “lag period” where no *Calluna* plant is present for a specified number of years after death or management, as was adopted by Allen et al., (2013). The model uses mortality data from Sike Hill in Moor House National Nature Reserve at an altitude of 550m. Growth rates are known to be dependent on temperature and altitude and therefore these establishment periods may be longer in more climatically challenging conditions (Armstrong et al., 1997). Grazing may also influence mortality, particularly if the site is overgrazed (Moss 1989). To understand the sensitivity of our results to these input data, we also analysed mortality rates between 25% lower and 25% higher than those observed by Forrest (1971) in the main analysis, and much wider ranges in the supplementary sensitivity analysis.

Assumption (d) does not hold on very long time scales (Gimingham, 1988; Marrs, 1986) because of factors such as catastrophic events and long-term changes brought about by succession, climate change, nutrient deposition, heather beetle (*Lochmaea suturalis*) or altered management. This is particularly true of *Calluna* heaths, however our model concerns blanket bog as this is the habitat where regulation has been updated to ban managed burning in the UK.

Multi-century periods of *Calluna* dominance at UK blanket bog sites has been shown in the palaeoecological record (Chambers 1982, Chamber 1983, Chambers 2022, Webb et al. 2022, McCarroll et al. 2016) so the conditions in our model are not without precedent. On timescales less than the wildfire return period (∼150 years average in the UK (Davies et al., 2016)), it is plausible that many stands of *Calluna* will persist for the timescale of our model, especially if grazing limits succession to a treed landscape, as is often the case in UK uplands (Moss 1989). Furthermore, our model’s purpose is not to predict accurately what a specific *Calluna* stand may look like at any given moment in time, but rather to estimate the time taken for a stand released from management to achieve a stable genuinely ‘unmanaged’ condition in the absence of other ecological disturbances. In reality, some stands will never actually reach this point due to the potential for vegetative change or disturbance brought about by other factors. We ran our model for double the average wildfire return period to understand how persistent management or wildfire effects were in this context. Although shorter periods would have more ecological grounding as succession would be much less likely, our purpose was to understand if management or wildfire effects would still persist before the next likely incidence of wildfire.

Our model takes a stand of 10,000 plants and ages each plant by a year at each time step. At each annual time step, every plant in the stand has an age-dependent chance of dying based on the data from Forrest (1971) by fitting an exponential relationship between age and mortality (Eq.1). When a plant is one year old it has a 0.99% chance of dying, compared with a 100% chance by age 44 in the central scenario with a mortality multiplier used to increase or decrease this rate in the sensitivity analysis. If the plant dies in a given year, the age counter is reset to zero (unless a lag period is specified). If the plant survives, it ages one year and the process repeats. Where a lag in establishment is used, the age counter is reset to -4 (five-year lag period accounting for 0), representing a gap in the *Calluna* cover until a positive number is reached.

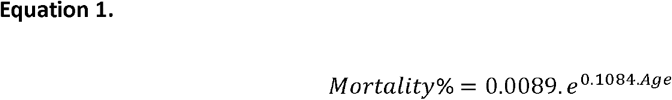

*Calluna* management was simulated in the model by resetting a proportion of the *Calluna* ages to zero value (or -4 when accounting for a lag period), with the “burn amount” defining what proportion was reset to zero and the “burn frequency” specifying at what frequency this occurred (e.g. 6.66% reset to zero each year to simulate a 15-year prescribed burn return period). The simulated stand would be divided into groups based on: number of plants / burn amount. The model cycled through which group of plants would be reset at each time step, such that each plant would be ‘burned’ once every 15 years, or whatever return period was specified.

We considered the time taken for a stand of *Calluna* to reach a stable distribution, either because no management was applied or because management had ceased. At this point we consider the stand to be of a mixed age as it no longer has an artificially narrow age distribution (Fig 1 A shows what we term an even aged distribution which is artificially narrow, Fig 1 B shows a mixed age stand in a more natural condition). We assessed this by analysing the age distribution of each year using the Kolmogorov-Smirnov test and comparing this to the mean age distribution in the final 10 years of the model run (years 291 to 300) for each scenario. Highly similar age distributions have a low Kolmogorov’s D statistic, whilst different age distributions have a high D statistic. P values are also calculated, indicating statistically significant differences between age distributions. Thus, a similar age distribution would have a low D statistic and high p value, while a statistically significant different age distribution would have a high D statistic and low p value. P values were adjusted using the “BY” method for multiple testing correction (Benjamini & Yekutieli, 2001). Other methods are explored in the supplementary sensitivity analysis but were found to give broadly similar results.

**Figure 1:**
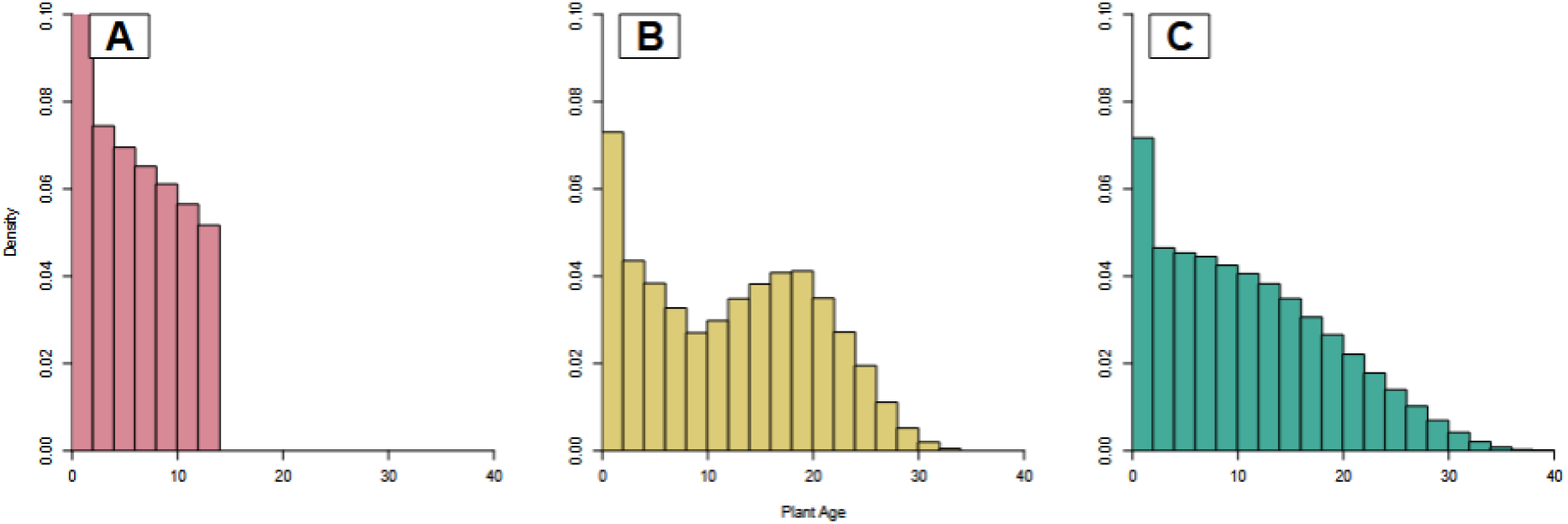
Age distribution of modelled *Calluna* plants shown as A, in a 15-year management cycle, B, in post-management disturbance, and C, having reached a stable final distribution.

We define a stable age distribution as the adjusted p value being less than 0.05 and for 10 consecutive years or more to ensure such a distribution is not transitory. This method gave similar results to using the D statistic, assessing statistically similar distributions using p-values with various types of corrections to control inflation of errors for multiple comparisons (Supplementary Figure 1).

We defined a series of scenarios to assess ways in which *Calluna* reaches a stable distribution (see Table 1). In the first scenario, no management is applied and we assessed how long it takes to reach a stable distribution. This first scenario would be analogous to a large area recovering from wildfire or other catastrophic event which affects all plants in the stand at the same time. In scenarios 2 to 4 we ran the model starting from the end distribution in scenario 1 and applying management return periods of 15, 20 and 25 years until a new stable distribution was reached. We then ceased management and assessed how long it takes to achieve a new stable distribution.

**Table 1:** Model parameters for the different scenarios.

| Scenario | Time<br>period<br>(years) | Start age | Management | Burn<br>start<br>year | Burn<br>end<br>year | Burn<br>frequency | Burn<br>amount<br>(%) |
| --- | --- | --- | --- | --- | --- | --- | --- |
| 1 | 300 | 1 | No | n/a | n/a | - | 0 |
| 2 | 300 | natural | Yes | 1 | 100 | Annually | 6.66 |
| 3 | 300 | natural | Yes | 1 | 100 | Annually | 5 |
| 4 | 300 | natural | Yes | 1 | 100 | Annually | 4 |

## Results

Our results for four scenarios, with both increased and lowered mortality rates taken from Forrest (1971), are presented in Table 2. These results use the five-year lag period suggested by Allen et al., (2013). The same scenarios without this lag period, which generally show lower times to reach a stable state, are available in Supplementary Information. We give the time taken for the model to reach a stable age distribution after the management/wildfire disturbance. We take this to be the time over which management/wildfire effects persist.

**Table 2:** Results of model analysis for scenarios with a five-year lag time for plant recovery, showing the number of years to reach the stable age distribution stage (based on all 1,000 model runs).

| Scenario | Mortality multiplier | Time to reach stable age distribution |
| --- | --- | --- |
| 1 | 0.75 | 88 ± 8 |
| 1 | 1 | 76 ± 7 |
| 1 | 1.25 | 67 ± 6 |
| 2 | 0.75 | 79 ± 8 |
| 2 | 1 | 66 ± 7 |
| 2 | 1.25 | 57 ± 7 |
| 3 | 0.75 | $68 \pm 8$ |
| 3 | 1 | $56 \pm 8$ |
| 3 | 1.25 | $47 \pm 7$ |
| 4 | 0.75 | $56 \pm 9$ |
| 4 | 1 | $44 \pm 7$ |
| 4 | 1.25 | $37 \pm 6$ |

A typical age distribution for scenario 2 during each stage is shown in Figure 1 which highlights a truncated, even-aged distribution in Fig 1 A brought about artificially by management. During the management stage, the plants reach a maximum age of 14, reflective of the 15-year complete burn cycle. Once released from management the distribution tends towards a wider, uneven age distribution reflecting a range of growth stages of the plants.

We show the change in similarity to an unmanaged age distribution in Figure 2 using the D statistic and p values over time (see Supplementary Information for comparison of statistical techniques). For scenario 1 (no management) we can see a long period of transition as plants naturally age, die and are replaced. For scenarios 2 to 4 there is a rapid shift to a new, stable age distribution under such management. Once management ceases, the time to reach a natural stable state is dependent on the return period of the previous management, with shorter return periods leading to longer post-management effects. See Supplementary Information for a sensitivity analysis of the mortality rate and burn percentage on our results.

**Figure 2:**
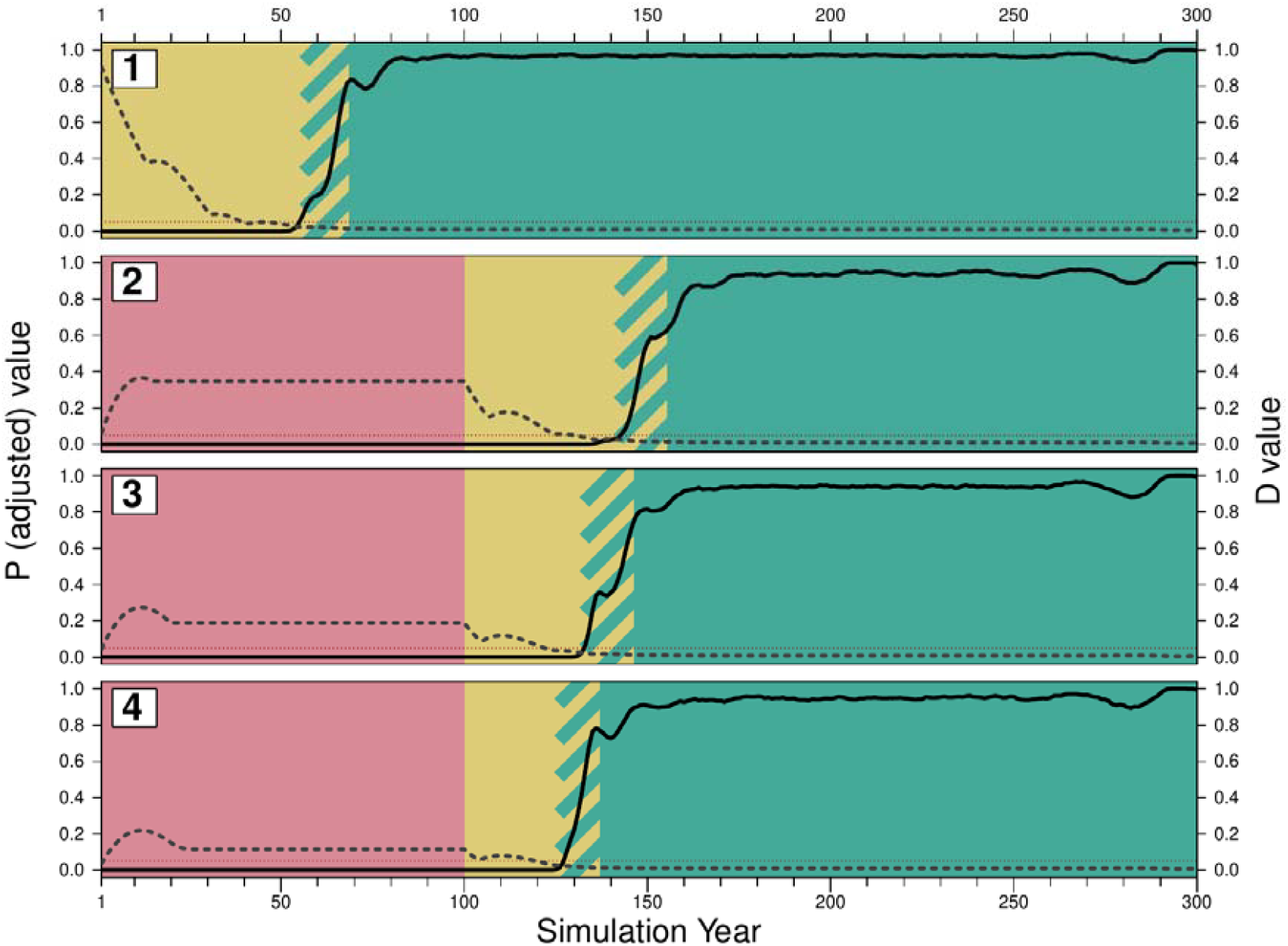
Similarity in age distribution of *Calluna* plants to an unmanaged condition over time during each of the model scenarios. The green area represents a stable, unmanaged distribution. The red area represents a period under management and the yellow area represents the transition. The hatched area indicates the standard deviation between the model runs for the start of the final stable period. Mean adjusted p values are shown as a solid black line, while d values are shown as a dashed grey line.

## Discussion

Our model addresses our initial aim a), to estimate the time taken to reach an unmanaged state after both catastrophic wildfire and the cessation of management. Considering the central mortality estimate and five-year lag time, our model suggests that it probably takes around 75 years for a *Calluna* stand to be free from post-disturbance effects on age distribution after a stochastic and catastrophic event such as wildfire where all plants are burned (Table 2, Scenario 1). If a previously managed stand is released from management, our model shows management effects will persist for a period dependent on the return period of the previous management. This ranges from approximately 62 years for a management return period of 15-years to around 44 years for a return period of 25 years (Table 2, Scenarios 2 and 4). This is the time taken to reach an age distribution which could be described as similar to an unmanaged stand. The time required to reach a stable ‘unmanaged’ age is dependent on mortality and the lag in *Calluna* establishment, which are both probably greater in climatically marginal areas where growth rates are slower (Armstrong et al., 1997), though they may also be influenced by grazing intensity (Moss, 1989). We address this issue further in the supplementary sensitivity analysis considering the effect of mortality rates on our different management scenarios. More climatically favourable sites will have lower mortality rates and consequently take a longer time to reach an uneven aged stand as individual plants are replaced at a slower rate meaning the management disturbance effect (plants of the same age) lasts longer.

Our model does not address changes in grazing intensity that may alter the establishment and mortality of plants, nor how changing climate may alter mortality rates if high altitude sites become less climatically marginal or water tables become lower at blanket bog sites both of which are possible giving shifting climatic envelopes for blanket bog (Ritson et al. 2025). The relative proportion of vegetative regrowth versus seedling establishment can also have implications for *Calluna* population dynamics (Scandrett and Gimingham 1989) and is not covered in our model. Similarly, the age of the *Calluna* stand and the intensity of the fire itself can also effect re-establishment of *Calluna* (Davise et al. 2010) and is not covered in our model. All of these issues would need to be addressed to accurately model future fuel loads, however this was not the purpose of our model. We instead adopt a more simplistic approach to understand the persistence of post-disturbance effects.

In the context of recent decisions to ban burning as a management tool on peat >30-40 cm deep in the UK, our model results provide insight into how long after the cessation of management a *Calluna* dominated blanket bog would require in order to return to a more mixed aged condition in which the competitive advantage of *Calluna* is reduced. During this period, fuel loads and severe wildfire risk would rise (Allen et al., 2013), although this would not be a simple increase because there is a non-linear relationship between annual biomass production and age (Forrest, 1971). The level of increased risk might be unacceptable given the predicted increase in wildfire frequency in the UK under climate change (Little et al., 2025), combined with the potential for intense wildfires under high fuel loads to ignite the peat itself, with catastrophic loss of carbon and habitat (Maltby et al., 1990). Our model suggests that, if all management were to be ended under a ‘rewilding’ scenario, it could take around 50 years until *Calluna* mortality creates a steady-state, mixed aged stand, depending on mortality and other factors. During this period fuel loads would be elevated above current, managed levels.

Alternative methods of decreasing *Calluna* standing biomass are therefore desirable if prescribed burning is no longer an option. The rewetting of bogs and the re-establishment of a *Sphagnum* understory can force *Calluna* into an alternate growth form known as ‘layering’ (Forrest 1971; Keatinge 1975; MacDonald *et al*., 1995) in which the growth of *Calluna* is constrained by the wet conditions and grows as small shoots through a *Sphagnum* layer. Even when a forced switch to the layering growth form is not achievable, wetter conditions generally still result in lower total aboveground biomass of *Calluna* because such conditions are sub-optimal for *Calluna* growth (Forrest 1971, Wallèn 1987). Even damaged or drained peatlands lead to lower biomass accumulation compared to *Calluna* growing in more optimal freely-draining heathland conditions (Grau-Andrés et al., 2018). Where wildfires do occur, the risk that these will develop into subsurface ‘smouldering’ fires which consume the peat is minimised if water levels are high (Prat-guitart et al., 2016).

It may be possible, therefore to decrease *Calluna* biomass accumulation and consequent risk and severity of wildfire through the use of peatland restoration and *Sphagnum* reintroduction, which has been successfully accelerated in trials (*e.g*. Caporn *et al*., 2018; Fandrem *et al*., 2025), although it remains to be seen to what extent this can be achieved everywhere, given climatic and direct human pressures on blanket bogs (Ritson et al. 2025). If layering within *Sphagnum*, or at least stunted *Calluna* growth through raised water tables, can be induced this has the added benefit of requiring minimal ongoing management once wet conditions have been established. Further work is needed to understand over what proportion of deep peat this could be achieved given factors such as slope, presence of gullies or artificial drainage networks, past management and atmospheric pollution, all of which may constrain development of suitable habitat for *Sphagnum* or the ability to raise water tables.

Mowing of *Calluna*, either to create firebreaks or to simulate the effects of prescribed burning of larger patches, is another potential management technique to limit severe wildfire risk (Defra 2025), but this has a number of disadvantages. Mowing, where used to mimic the effects of burning, maintains a narrow age structure in the *Calluna* stands, potentially limiting biodiversity (Watt 1955, Gimingham 1960). In the absence of rewetting, it also requires regular management, while regular use of machinery brings with it potential compression of the peat surface. Indeed a recent review has highlighted the need for more evidence exploring the impacts of mowing on peat because of concerns over the potential interplay of such factors (Moody and Holden, 2023). On the other hand, management through mowing may have the potential to enhance carbon capture into peatlands via re-application of harvested biomass, or conversion of the harvested material to biochar to enhance organic matter persistence either on-site or elsewhere (Jeewani et al., 2025).

Answering our initial aim b) concerning the validity of chronosequences of *Calluna* sites, our results also highlight a significant issue with much of the existing literature concerning *Calluna* management, which is that stands that have been burned relatively recently are often used as reference for an unburned or unmanaged condition. For example, Davies and Legg, (2008) use 25 to 30 years as being ‘long term unburned’, Harris et al. (2011) use >35 years as their cutoff for a moor to be classed as ‘unburned’, and Heinemeyer et al. (2023) use a value of 22-25 years post-burn as being ‘unmanaged’. Our results suggest that at least double that time would be needed to exclude the effects of previous management. Studies using time-periods less than ∼50 years as an ‘unmanaged’ reference for a *Calluna* community may therefore not have been studying a truly unmanaged condition but rather a system still in post-management recovery. Our model shows that the extent of any resulting (unintended) bias would likely depend on the age of *Calluna* and the time of observation in these studies. In general, however, we would expect all studies undertaken during this recovery period to over-estimate the risk of wildfire compared to that of a genuinely natural, or fully recovered, blanket bog. The use of sites with persistent management effects as natural reference sites has important consequences for our understanding of severe wildfire risk on peatlands. It has likely led to over-estimation of wildfire risks on natural blanket bogs and overly pessimistic assessments of the ‘restorability’ of historically burn-managed areas. While elevated severe wildfire risk during the post-management period is real, it is also in part transient and may be minimised by active interventions such as those described above.

Anthropogenic changes to fire frequency, pattern and intensity can have dramatic changes on ecological systems where fire management can be viewed as a perturbation as compared to the natural cycle of fire disturbance (Keeley and Pausas., 2019). Our model highlights the difficulty in finding genuinely unmanaged analogues to compare against in these heavily anthropogenic habitats when fire frequency, patterning and intensity are all known to have changed in the last ∼100 years and have likely interacted with other management and climatic pressures (Ritson and Lindsay 2023). This work sounds a note of caution for any study using a site as an unmanaged analogue without first understanding site history and the likely persistence of any previous management or disturbance.

## Conclusions

Our model shows that cessation of management will have effects for around 50 years, persisting until *Calluna*-dominated areas reach a new stable age distribution. During this time fuel loads will be elevated as the narrow age distribution of plants created by management ages. Given the elevated risk of severe wildfire brought about by this increased fuel load, coupled with local and climatic pressures, some form of active management is likely needed to mitigate this risk or accelerate a transition to a less *Calluna* dominated system.. Given that managed burning is to be banned on peat >30-40 cm deep in the UK, such mitigating measures could either take the form of limiting *Calluna* growth through rewetting and Sphagnum reintroduction, or of mowing the *Calluna*. The efficacy of these interventions in delivering reduced fuel loads, adequate forage and nesting for grouse and other ecosystem services from blanket bogs still needs to be assessed. Our results have wider relevance for any managed systems in that they highlight the challenge of finding truly unmanaged analogues to compare management effects against due to the often underestimated persistence management.

## Supporting information

Supplementary sensitivity analysis

## Acknowledgements

All authors were funded by the UKRI Greenhouse Gas Removals Demonstrator – Peat project (Grant Number BB/V011561/1).

## References

Allen, K.A., Harris, M.P.K., Marrs, R.H., 2013. Matrix modelling of prescribed burning in Calluna vulgaris-dominated moorland: Short burning rotations minimize carbon loss at increased wildfire frequencies. J. Appl. Ecol. 50, 614–624. 10.1111/1365-2664.12075

Armstrong, H.M., Gordon, I.J., Grant, S.A., Hutchings, N.J., Milne, J.A., Sibbald, A.R., 1997. A Model of the Grazing of Hill Vegetation by the Sheep in the UK. I. The Prediction of Vegetation Biomass. J. Appl. Ecol. 34, 166. 10.2307/2404857

Barclay-Estrup, P., Gimingham, C.H., 1969. The Description and Interpretation of Cyclical Processes in a Heath Community: I. Vegetational Change in Relation to the Calluna Cycle. J. Ecol. 57, 737. 10.2307/2258496

Benjamini, Y., Yekutieli, D., 2001. The control of the false discovery rare in multiple testing under dependency. Ann. Stat. 29, 1165–1188.

Caporn, S., Rosenburgh, A., Keightley, A., Hinde, S., Riggs, J., Buckler, M., & Wright, N. (2018). Sphagnum restoration on degraded blanket and raised bogs in the UK using micropropagated source material: a review of progress. Mires and Peat, 20, 1–17. 10.19189/map.2017.omb.306

Chambers, F.M., 1982. Two radiocarbon-dated pollen diagrams from high-altitude blanket peats in South Wales. The Journal of Ecology, pp.445–459.

Chambers, F.M., 1983. Three radiocarbon-dated pollen diagrams from upland peats north-west of Merthyr Tydfil, South Wales. The Journal of Ecology, pp.475–487.

Chambers, F. (2022) The use of paleoecological data in mire and moorland conservation. PAGES, 30(1), 16–17

Matt Davies, G., Adam Smith, A., MacDonald, A.J., Bakker, J.D. and Legg, C.J., 2010. Fire intensity, fire severity and ecosystem response in heathlands: factors affecting the regeneration of Calluna vulgaris. Journal of Applied Ecology, 47(2), pp.356–365.

Davies, G.M., Kettridge, N., Stoof, C.R., Gray, A., Ascoli, D., Fernandes, P.M., Marrs, R., Allen, K.A., Doerr, S.H., Clay, G.D., McMorrow, J., Vandvik, V., 2016. The role of fire in UK peatland and moorland management: The need for informed, unbiased debate. Philos. Trans. R. Soc. B Biol. Sci. 371. 10.1098/rstb.2015.0342

Davies, G.M., Legg, C.J., 2008. The effect of traditional management burning on lichen diversity. Appl. Veg. Sci. 11, 529–538. 10.3170/2008-7-18566

Defra, 2007. The Heather and Grass Burning Code. London, UK.

Defra 2025. Heather and Grass Management Code 2025, London, UK.

Fandrem, M., Hassel, K. and Speed, J.D.M., Kolstad, A.L and Kyrkjeeide, M.O. (2025) From bare peat to Sphagnum cover: The success of Sphagnum-fragment and straw-mulch application for initiating peatland restoration. Mires and Peat, 32, Article 05. DOI: 10.19189/001c.131833

Forrest, G., 1971. Structure and Production of North Pennine Blanket Bog Vegetation. J. Ecol. 59, 453–479.

Gimingham, C.H. (1960) Biological Flora of the British Isles. Calluna vulgaris (L.) Hull. J. Ecol, 48, 455–483.

Gimingham, C.H., 1988. A reappraisal of cyclical processes in Calluna heath. Vegetatio 77, 61–64. 10.1007/BF00045751

Gimingham, C.H., 1972. The ecology of heathlands. Chapman and Hall, UK.

Harper, A.R., Doerr, S.H., Santin, C., Froyd, C.A., Sinnadurai, P., 2018. Prescribed fire and its impacts on ecosystem services in the UK. Sci. Total Environ. 624, 691–703.

Holden, J., Shotbolt, L., Bonn, A., Burt, T.P., Chapman, P.J., Dougill, A.J., Fraser, E.D.G., Hubacek, K., Irvine, B., Kirkby, M.J., Reed, M.S., Prell, C., Stagl, S., Stringer, L.C., Turner, A. and Worrall, F. (2007) Environmental change in moorland landscapes. Earth Science Reviews, 82, 75–100.

Jeewani, P.H., Brown, R.W., Evans, C.D., Cook, J., Roberts, B.P., Fraser, M.D., Chadwick, D.R. and Jones, D.L., 2025. Rewetting alongside biochar and sulphate addition mitigates greenhouse gas emissions and retain carbon in degraded upland peatlands. Soil Biology and Biochemistry, 207, p.109814.

Keeley, J. E., & Pausas, J. G. (2019). Distinguishing disturbance from perturbations in fire-prone ecosystems. International Journal of Wildland Fire, 28, 282–287.

Leslie, A.., 1911. The grouse in health and in disease[]: being the popular edition of the report of the committee of inquiry on grouse disease. Smith Elder and Co, London, UK.

Little, K., Vitali, R., Belcher, C.M., Kettridge, N., Pellegrini, A.F.A., Ford, A.E.S., Smith, A.M.S., Elliott, A., Voulgarakis, A., Stoof, C.R., Kolden, C.A., Schwilk, D.W., Kennedy, E.B., Newman Thacker, F.E., Millin-Chalabi, G.R., Clay, G.D., Morison, J.I., McCarty, J.L., Ivison, K., Tansey, K., Simpson, K.J., Jones, M.W., Mack, M.C., Fulé, P.Z., Gazzard, R., Harrison, S.P., New, S., Page, S.E., Hall, T.E., Brown, T., Jolly, W.M., Doerr, S., 2025. Priority research directions for wildfire science: Views from a historically fire-prone and an emerging fire-prone country. Philos. Trans. R. Soc. B Biol. Sci. 380. 10.1098/rstb.2024.0001

Maltby, E., Legg, C.J., Proctor, M.C.F., 1990. The Ecology of Severe Moorland Fire on the North York Moors: Effects of the 1976 Fires, and Subsequent Surface and Vegetation Development. J. Ecol. 78, 490. 10.2307/2261126

Marrs, R.H., 1986. The role of catastrophic death of Calluna in heathland dynamics. Vegetatio 66, 109–115. 10.1007/BF00045500

McCarroll, J., Chambers, F.M., Webb, J.C. and Thom, T., 2016. Using palaeoecology to advise peatland conservation: An example from West Arkengarthdale, Yorkshire, UK. Journal for Nature Conservation, 30, pp.90–102.

Moody, C.S., Holden, J., 2023. The Impacts of Vegetation Cutting on Peatlands and Heathlands - Natural England Evidence Review NEER028. York, UK.

Moss, C.E. (1904) Peat moors of the Pennines: their age, origin and utilisation, The Geographical Journal, 23(5), 660–671.

Moss, R., 1989. Management of heather for game and livestock. Bot. J. Linn. Soc. 191, 301–306.

Noble, A., Palmer, S.M., Glaves, D.J., Crowle, A., Holden, J., 2019. Peatland vegetation change and establishment of re-introduced Sphagnum moss after prescribed burning. Biodivers. Conserv. 28, 939–952. 10.1007/s10531-019-01703-0

Noble, A., Glaves, D.J., Leppitt, P., Crowle, A., Key, D. and Rodgers, A. 2025. An evidence review update on the effects of managed burning on upland peatland biodiversity, carbon and water. Natural England Evidence Review, NEER155. Natural England.

Prat-guitart, N., Rein, G., Hadden, R.M., Belcher, C.M., Yearsley, J.M., 2016. Effects of spatial heterogeneity in moisture content on the horizontal spread of peat fi res. Sci. Total Environ. 572, 1422–1430. 10.1016/j.scitotenv.2016.02.145

R Core Team (2022). R: A language and environment for statistical computing. R Foundation for Statistical Computing, Vienna, Austria. https://www.R-project.org/.

Rennie, G., Brown, R., Shirreff, J. (1794) General View of the Agriculture of the West Riding of Yorkshire: With Observations on the Means of its Improvement. W. Bulmer and Company, London, UK.

Ritson, J.P., Lindsay, R.A., 2023. The use of historical accounts of species distribution to suggest restoration targets for UK upland mires within a ‘moorland’ landscape. Mires Peat 29, 1–17.10.19189/MaP.2023.OMB.Sc.2116686

Ritson, J. P., Lees, K. J., Hill, J., Gallego-Sala, A., & Bebber, D. P. (2025). Climate change impacts on blanket peatland in Great Britain. J. Appl. Ecol. December 2024, 1–14. 10.1111/1365-2664.14864

Scandrett, E. and Gimingham, C.H., 1989. A model of Calluna population dynamics; the effects of varying seed and vegetative regeneration. Vegetatio, 84(2), pp.143–152.

Wallèn, B. (1987) Growth Pattern and Distribution of Biomass of Calluna vulgaris on an Ombrotrophic Peat Bog. Holarctic Ecology, 10(1), 73–79.

Watt, A.S., 1955. Bracken Versus Heather, A Study in Plant Sociology. J. Ecol. 43, 490–506.

Webb, J.C., McCarroll, J., Chamber, F.M., Thom, T. (2022) Evidence for the Little Ice Age in upland northwestern Europe: Multiproxy climate data from three blanket mires in northern England. The Holocene, 32(5), 451–467

