## Supplementary sensitivity analysis for "Modelling the persistence of post-management disturbance in *Calluna vulgaris* communities"

**Supplementary Information**

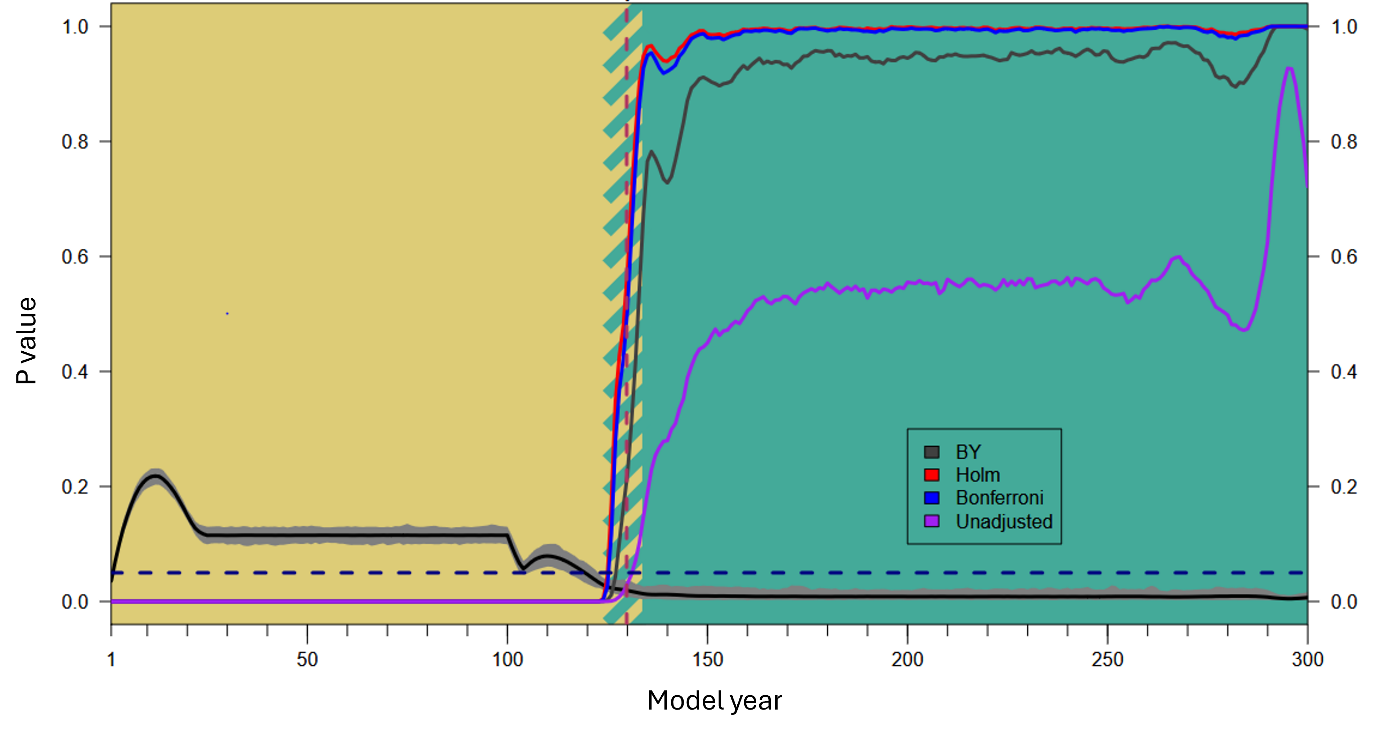

**Figure S1**: Comparison of different p-value corrections for multiple comparison for scenario 4. The blue dashed line indicated significant p values. The red dashed line is the mean value of detected difference in distribution from this method with the hatched area representing the standard deviation of this value. As all methods fall within the standard deviation of our chosen method, we assume our results are largely indifferent to the choice of statistical method.

**Table S1.** Results of model analysis for scenarios with no lag time for plant recovery, showing the number of years to reach the stable age distribution (based on all 1,000 model runs).

| **Scenario** | **Mortality multiplier** | **Time to reach stable age** |
| --- | --- | --- |
| 1 | 0.75 | 74 ± 6 |
| 1 | 1 | 63 ± 6 |
| 1 | 1.25 | 56 ± 5 |
| 2 | 0.75 | 60 ± 7 |
| 2 | 1 | 50 ± 6 |
| 2 | 1.25 | 42 ± 5 |
| 3 | 0.75 | 50 ± 7 |
| 3 | 1 | 40 ± 6 |
| 3 | 1.25 | 40 ± 5 |
| 4 | 0.75 | 40 ± 7 |
| 4 | 1 | 32 ± 5 |
| 4 | 1.25 | 28 ± 4 |

**Supplementary sensitivity analysis**

**Scenario 1 – no management**

We assessed the effect of the mortality multiplier on the time taken to reach a stable age distribution. We performed 1000 model runs for each scenario varying the mortality rate and calculated means and standard deviations for each batch of model runs. We then used linear regression to assess the relationship between the two variables. Significant negative correlations were noted for all scenarios (r=0.99, p<0.001 for all tests). This modelling was performed for scenarios with no lag period; a five-year lag period; and a ten-year lag period (Figure S1). Significant negative correlations were noted for all scenarios with the following equations

**0 lag**

y = 66.406 - 29.869 * log_10_(mortality multiplier)

**5-year lag**

y = 78.394 - l34.731 * log_10_(mortality multiplier)

**10-year lag**

y = 100.334 - 39.960 * log_10_(mortality multiplier)
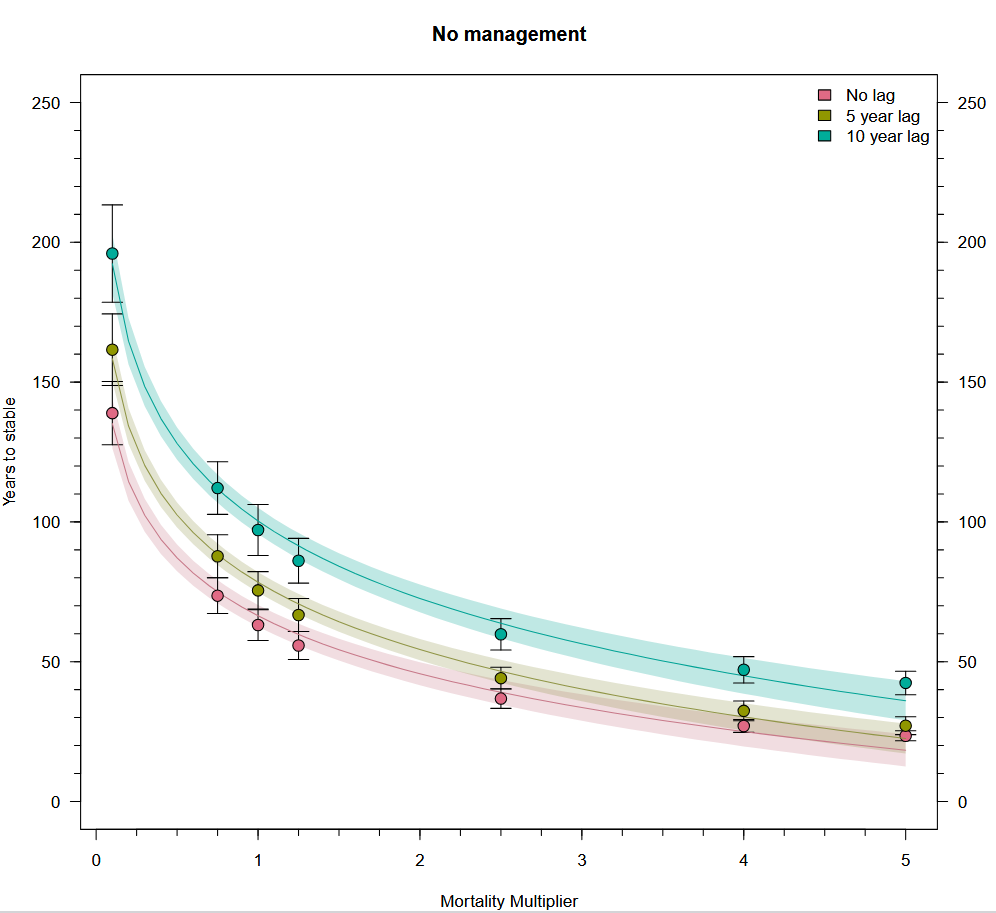

Figure S2: relationship between time taken to reach a stable state and the mortality multiplier for different lag periods for Calluna establishment (error bars at one standard deviation of the 1,000 model runs).

**Scenario 4 – Management starting from a stable age distribution**

We then assessed the impact of annual burn % on the time take to reach a stable age distribution in a management scenario. We performed 1000 model runs for each scenario varying the burn % and calculated means and standard deviations for each batch of model runs (Figure S2). We then used linear regression to assess the relationship between the two variables. There was a significant negative correlation between time to reach stable age distribution and burn % (r=0.90, p=0.001) with an equation of:

**0 lag**

y = 21.054 + 12.663 * log_10_(burn %).

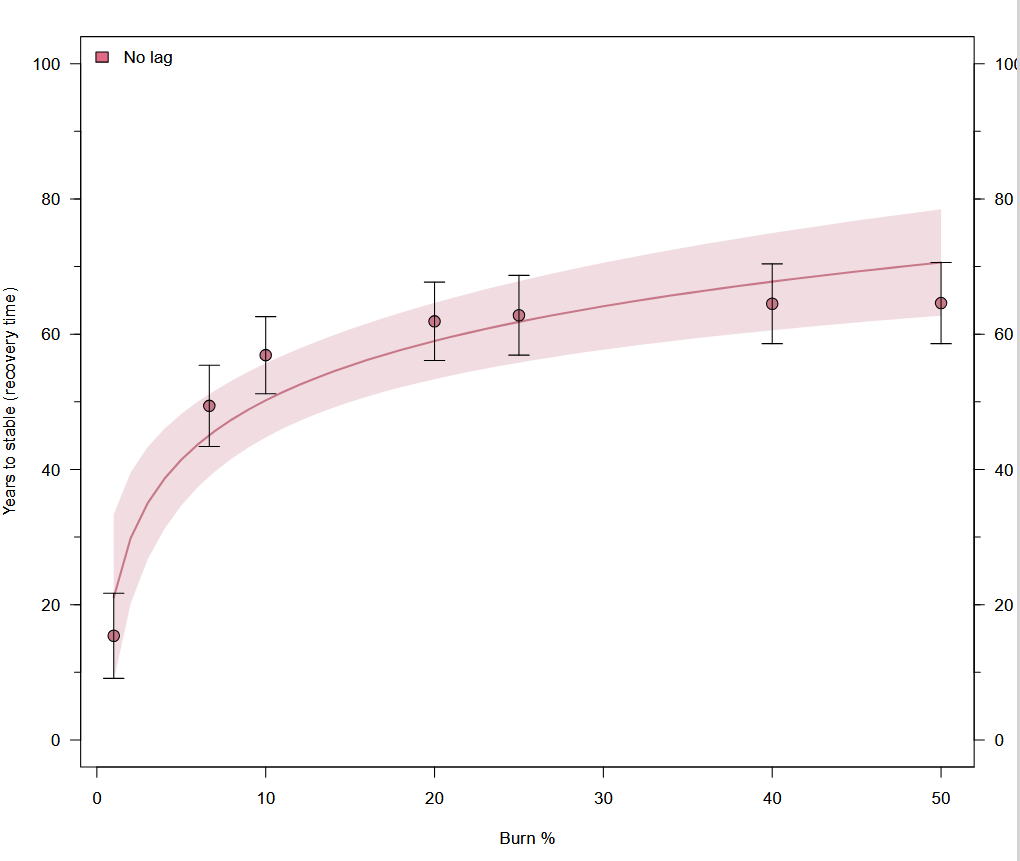

Figure S3: Relationship between time to stable age distribution and annual burn % during management (error bars at one standard deviation of the 1,000 model runs).
